# APOE Impacts Lipid Trafficking in Retinal Pigment Epithelium Cells

**DOI:** 10.1101/2024.05.30.596647

**Authors:** Sarah E.V. Richards, John Demirs, Sandra Jose, Lin Fan, YongYao Xu, Robert Esterberg, Chia-Ling Huang, Christopher W. Wilson, Magali Saint-Geniez, Sha-Mei Liao

## Abstract

Age-related macular degeneration (AMD) is typified by the formation of lipid-rich drusen under the retinal pigment epithelium (RPE) layer. Apolipoprotein E (APOE) is a known genetic risk factor for AMD and a substantial component of drusen, however, the mechanism by which APOE variants contribute to AMD pathology remains unclear. APOE is the primary cholesterol and lipid transport protein of the central nervous system, as well as a component circulating lipoproteins. To better understand how APOE-dependent lipid transport may impact AMD risk, we generated isogenic APOE iPS-RPE cells expressing each of the common human APOE isoforms, as well as an APOE knockout line. APOE knockout cells showed significant morphological and barrier function deficits, suggesting that APOE is essential for RPE health. Furthermore, we observed that APOE abundance is isoform-dependent in RPE cells and that lipid transport is deficient in APOE knockout RPE cells, as well as in RPE cells expressing APOE2, a variant associated with higher risk of AMD. Contrastingly, cells expressing APOE4 seem to respond strongly to lipid challenges by upregulating APOE to support efficient lipid transport. Our results suggest that disease associated APOE variants may impact lipid transport in RPE, contributing to the formation of drusen and impairing cellular function.

## Introduction

Age-related macular degeneration (AMD) is a leading cause of blindness in older adults^1–3^. AMD disease progression follows two primary trajectories, with both leading to progressive vision loss and possible blindness. Neovascular or “wet” AMD is typified by the growth of weak, leaky choroidal blood vessels whereas dry AMD results from progressive retinal pigment epithelium (RPE) dysfunction prior to overt regional RPE loss, termed geographic atrophy (GA). While age itself is the greatest risk factor for both forms of AMD, several protective and risk genetic variants have recently been identified^4–8^. Among these variants is apolipoprotein E (APOE), a cholesterol and lipid transport protein found on lipoproteins in both the systemic circulation and those produced locally in the central nervous system (CNS), including the retina. The three common human *APOE* isoforms, *APOE2*, *APOE3*, and *APOE4*, constitute an allelic series of AMD risk. *APOE4* is consistently associated with reduced risk for AMD, whereas *APOE2* is associated with increased AMD risk compared to the neutral *APOE3*^9–13^. *APOE4* is relatively common in most populations with an average allele frequency of about 20%, whereas *APOE2* accounts for only about 5% of alleles^14^, suggesting that understanding the role of *APOE* in AMD pathology has high relevance to a substantial portion of the population.

The role of *APOE* in AMD remains incompletely understood. However, years of intense study of *APOE* have revealed isoform-dependent differences that are conserved across biological systems and may be related to AMD. Compared to APOE3 or the protective APOE4, APOE2 shows profound deficits in binding to the low-density lipoprotein receptor (LDLR)^15–19^. Deficient LDLR binding results in increased abundance of extracellular APOE2 in the serum and most studied tissues due to a failure of LDLR-dependent APOE uptake and recycling. APOE isoforms also differ in lipid binding properties. Compared to APOE3, APOE4 preferentially binds larger lipoprotein particles such as LDL and VLDL, whereas APOE2 is more likely to bind HDL and small LDL particles^20^. High serum HDL concentrations are a known risk factor for AMD^21–23^, perhaps suggesting a link between *APOE* isoform, relative abundance of lipoprotein species, and AMD risk. Higher concentrations of secreted APOE could also lead to APOE accumulation in the extracellular space.

Previous reports indicate that APOE is a prevalent protein co-localized with drusen in AMD^24–29^. However, it is not known whether *APOE* isoform is correlated with drusen burden generally or if the *APOE2* risk allele is more likely to become incorporated into drusen. Despite some contradictory findings, work in the Alzheimer’s Disease field generally suggests that APOE4 binds less stably with amyloid-β^30–35^, perhaps suggesting that it has a lower propensity to accumulate in tissues in pro-aggregation conditions. However, it is unclear how this finding directly relates to the more heterogeneous protein and lipid aggregates forming drusen in AMD^24–29,36^. Interestingly, recent work has revealed *APOE* isoform-dependent differences in immunogenicity once APOE is deposited in plaques^9,37–39^, with APOE4 showing reduced immune activation and APOE2 increased activation in neural tissues. Previous work suggests that APOE4 is more likely to aggregate than the wildtype APOE3 ^40^, but a direct study of aggregation propensity of *APOE* isoforms in the context of AMD has not been explored. Recent work does suggest that APOE2 is more prone to forming stable intracellular aggregates in RPE cells that in turn disrupt cellular metabolism and induce RPE stress^41,42^. Substantial additional characterization is required to fully understand the role of *APOE* in AMD.

In the current work, we sought to understand how APOE functions in RPE using human iPSC-derived RPE cells (iPS-RPE) as an *in vitro* model system. Because APOE is implicated in many diverse functions in the brain and other organ systems, we focused only on those APOE isoform-dependent phenotypes that are typically conserved across systems. Our work establishes that APOE is required for RPE survival *in vitro*, and that APOE2 partially phenocopies total APOE loss of function (APOE KO) in some basic RPE functional assays. Similar to findings in other systems, we observe an increase in secreted APOE2 abundance compared to APOE3 or APOE4^43,44^. Consistent with a role of APOE in lipid trafficking and metabolism, bulk transcriptomics analysis revealed several lipid pathway disruptions resulting from loss of APOE in RPE cells. Importantly, we show that APOE4 iPS-RPE cells strongly upregulate APOE expression and increase lipid droplet storage in response to lipid challenge. Taken together, these results suggest that RPE-derived APOE shares similarities to APOE in other systems, with some notable differences in response to lipid challenge.

## Results

### APOE is abundant in soft drusen in human donor eye tissue

Previous reports have provided evidence that APOE is a common component in drusen, a hallmark of AMD pathology. Taken together with the known genetic risk for AMD associated with different APOE variants, inclusion of APOE in drusen is a key reason for interest in understanding the role of APOE in AMD pathology and progression. To better understand where APOE is found in AMD, we performed histology to visualize APOE as previously described^45^ in donor tissues from aged individuals without AMD pathology and with intermediate AMD (**Fig. 1**). APOE was present in normal aging donor samples in sparse cell populations in the neural retina and strongly enriched in the choroid (**Fig 1A**). Some normal aging samples also showed APOE accumulation in the Bruch’s membrane in the absence of any overlying pathology (**Fig 1A**, right). In intermediate AMD samples (**Fig 1B&C**), both hard and soft drusen were visible in the macula. Virtually all soft drusen stained strongly positive for APOE (**Fig 1B**), in addition to the choroid and in scattered cells of the neural retina. However, we rarely found hard drusen (**Fig 1C**) that were positive for APOE through the drusen mass. APOE was observed to accumulate on the outside of the hard drusen and in between hard drusen (**Fig 1C**, right), but not within the druse core. Previous reports suggest that soft drusen are enriched in lipids and lipoproteins compared to hard drusen, consistent with the high levels of APOE we observe in soft drusen^36,46^. These findings suggest that APOE protein accumulation is associated primarily with soft drusen, the more reliable drusen subtype for predicting AMD progression and severity.

**Figure 1:**
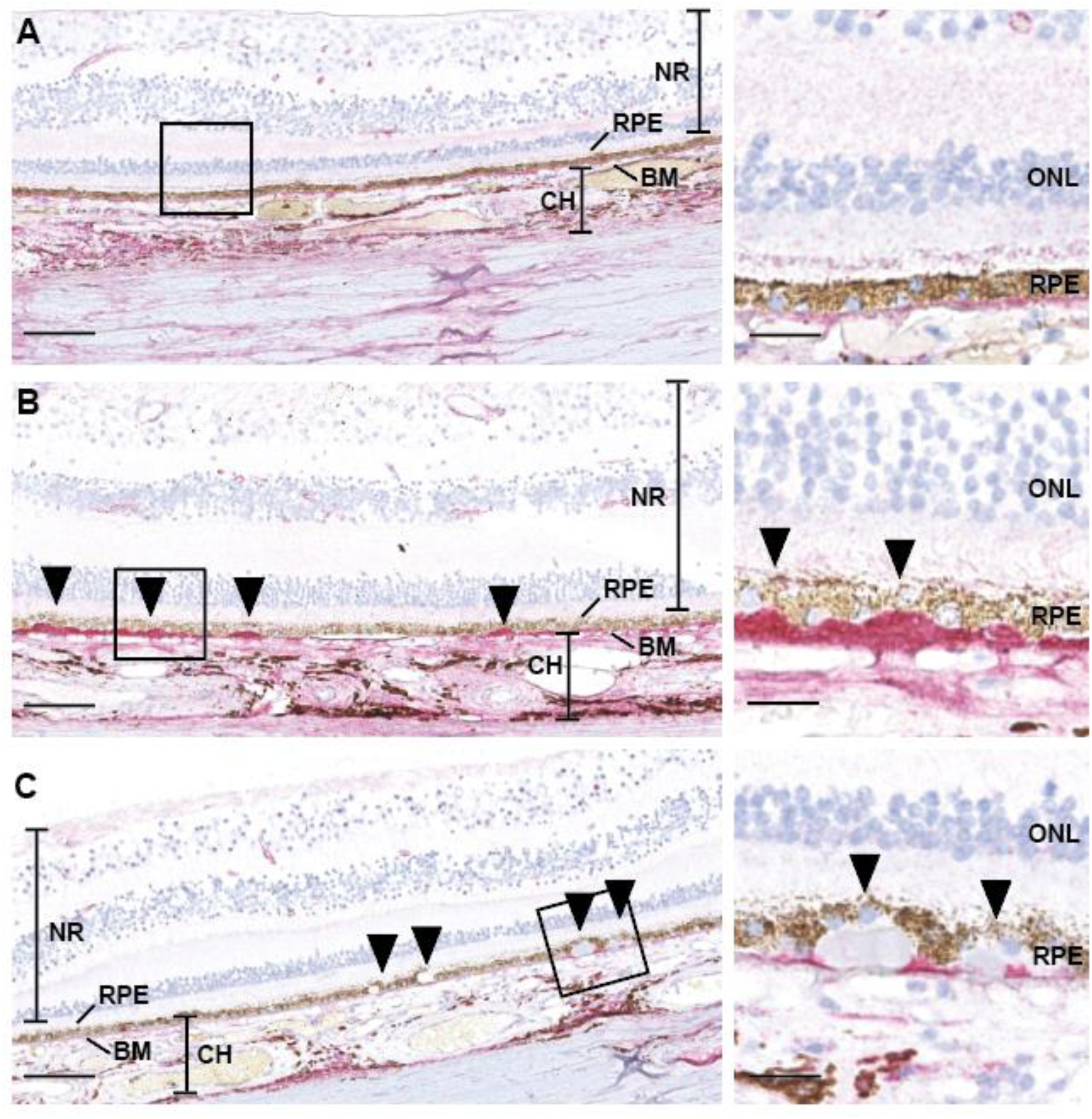
APOE accumulates in soft drusen, but rarely hard drusen in intermediate AMD eyes. **A)** Example of donor eye tissue from an aged individual without AMD pathology. APOE is present in the neural retina (NR), Bruch’s membrane (BM), and choriocapillaris (CH) but not in the Retinal Pigment Epithelium (RPE). **B)** Example of donor eye tissue from an aged individual with intermediate AMD pathology. Soft drusen underlying the RPE layer (black arrows) are strongly positive for APOE throughout their mass. **C)** Example of donor eye tissue from an aged individual with intermediate AMD pathology. APOE accumulates around, but not inside, hard drusen (black arrows), small drusen with clearly defined boarders. For all panels, area in square outline is the high magnification image in the right subpanel.

### Loss of APOE results in morphological and barrier function deficits in iPS-RPE cells

To better understand the function of RPE-derived APOE, iPS-RPEs expressing each of the common human *APOE* isoforms, as well as *APOE* knockout (KO) cells, were generated using CRISPR-Cas9 gene editing (**Fig 2A**). *APOE2*, *E3* and *E4* cells grew as expected, showing typical iPS-RPE hexagonal morphology and pigmentation throughout the 4-8weeks they were allowed to mature in culture on transwell inserts (**Fig 2B**). In contrast, APOE KO iPS-RPE appeared normal after initial seeding and matured for 2-3 weeks before beginning to show abnormal morphology and eventually dying in the center of each well and leaving behind only a ring of cells on the edge of each transwell insert (**Fig 2B**, **Fig S1A**). All genotypes were positive for RPE markers BEST1 (data not shown) and CRALBP (**Fig S1B**) and showed strong ZO-1 staining at cell boundaries (**Fig 2C**). Cell surface area was assessed using an automated image analysis pipeline to segment individual cells using phalloidin staining (**Fig 2D**). Both APOE2 and APOE KO cells had a larger average surface area than APOE3 and APOE4 cells (**Fig 2D**). However, APOE2 cells were uniformly larger in size while APOE KO cell populations were a heterogeneous mix of much smaller than average and much larger than average cells (**Fig S1C**). Transepithelial electrical resistance (TEER) was strongly impacted by APOE loss, with APOE KO cells having significantly lower TEER than all APOE-expressing iPS-RPE cells (**Fig 2E**).

**Figure 2:**
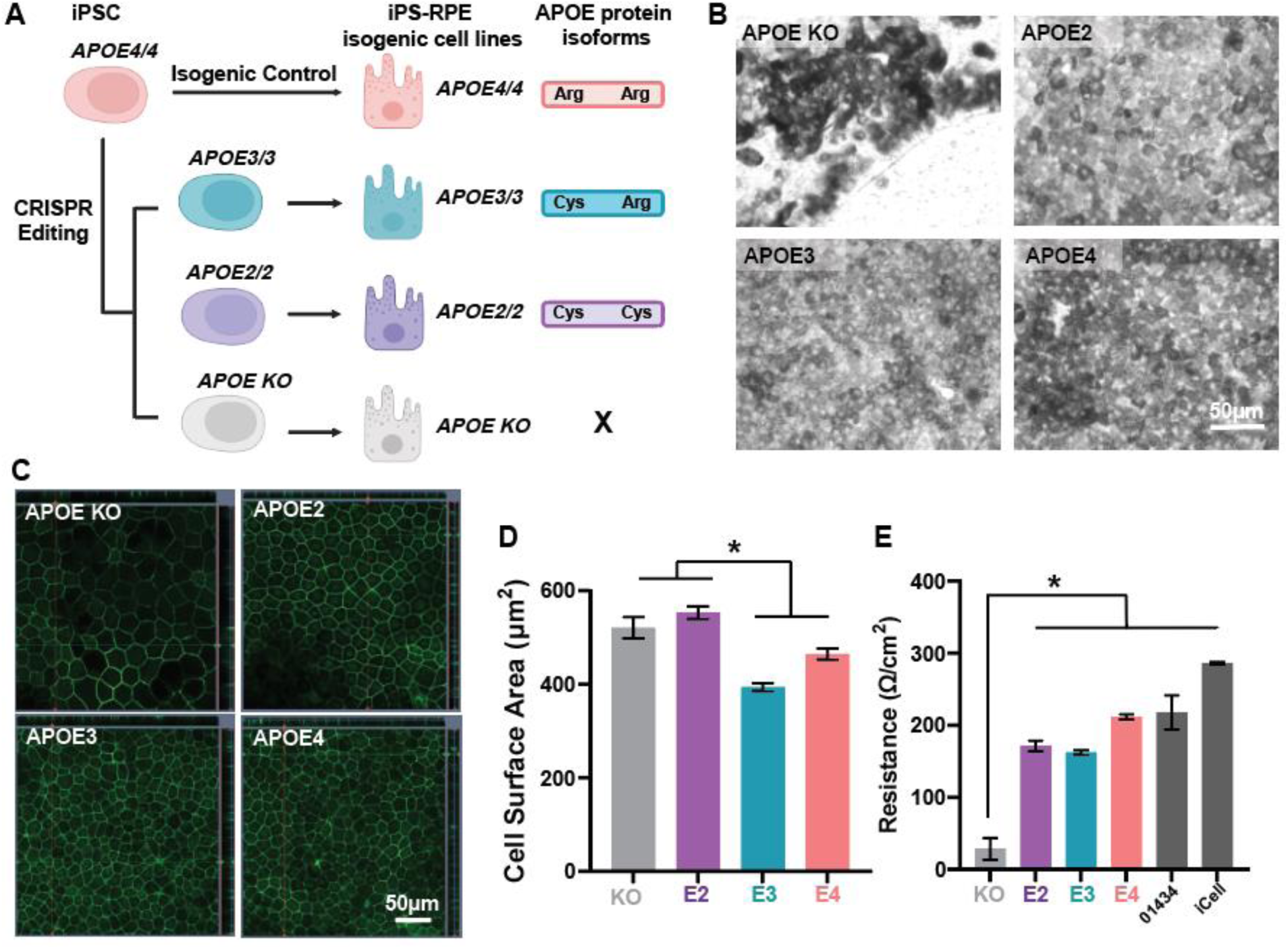
Loss of APOE is associated with severe morphological and barrier defects in iPS-RPE. **A)** Diagram of CRISPR editing strategy for APOE knockout and APOE isoform human iPSCs and iPS-RPE cells. **B)** Representative brightfield images of APOE iPS-RPE cells at 8 weeks in culture. Note severe pigment and morphological deficits in APOE KO (lower right) **C)** Representative images of ZO-1 staining in 8-week-old APOE iPS-RPE cells. Scale bar is 50µm. **D)** Cell surface area assessed from maximum intensity projections of phalloidin stained 8-week-old APOE iPS-RPE cells. **E)** Transepithelial electrical resistance (TEER) measured in 8-week-old APOE iPS-RPE cells and commonly used human iPS-RPE lines (01434 and FujiFilm iCell). * used to indicate any p <0.05 for simplicity.

### Secreted APOE2 is more abundant than APOE3 or APOE4

Previous reports suggest that APOE abundance varies by isoform both systemically and in the CNS^43,44^. APOE2 levels are generally found to be the highest of the three, with APOE4 being the lowest and APOE3 falling in the middle, perhaps resulting from deficient binding of APOE2 and LDLR, a major contributor to APOE endocytosis and recycling^19,47–49^. However, it is not known whether this is true for RPE cells or other ocular cell types. We measured APOE concentration using ELISA in media conditioned for 72 hours by iPS-RPE cells grown on transwell inserts. In all cells, we observed higher APOE abundance in the apical compartment than the basal compartment (**Fig 3A**). APOE2 cells showed the highest abundance of APOE in the apical compartment (**Fig 3A**), followed by APOE3 and then APOE4 (E2>>E3>E4) as we would expect from previous reports in other cell types and tissues. No APOE was detected in media samples from APOE KO cells (**Fig 3A**). Interestingly, *APOE* mRNA expression was not altered by *APOE* isoform (**Fig S1D**), suggesting that differences in secreted APOE abundance result from changes at the protein translation, degradation, or secretory levels, not at the level of transcription.

**Figure 3:**
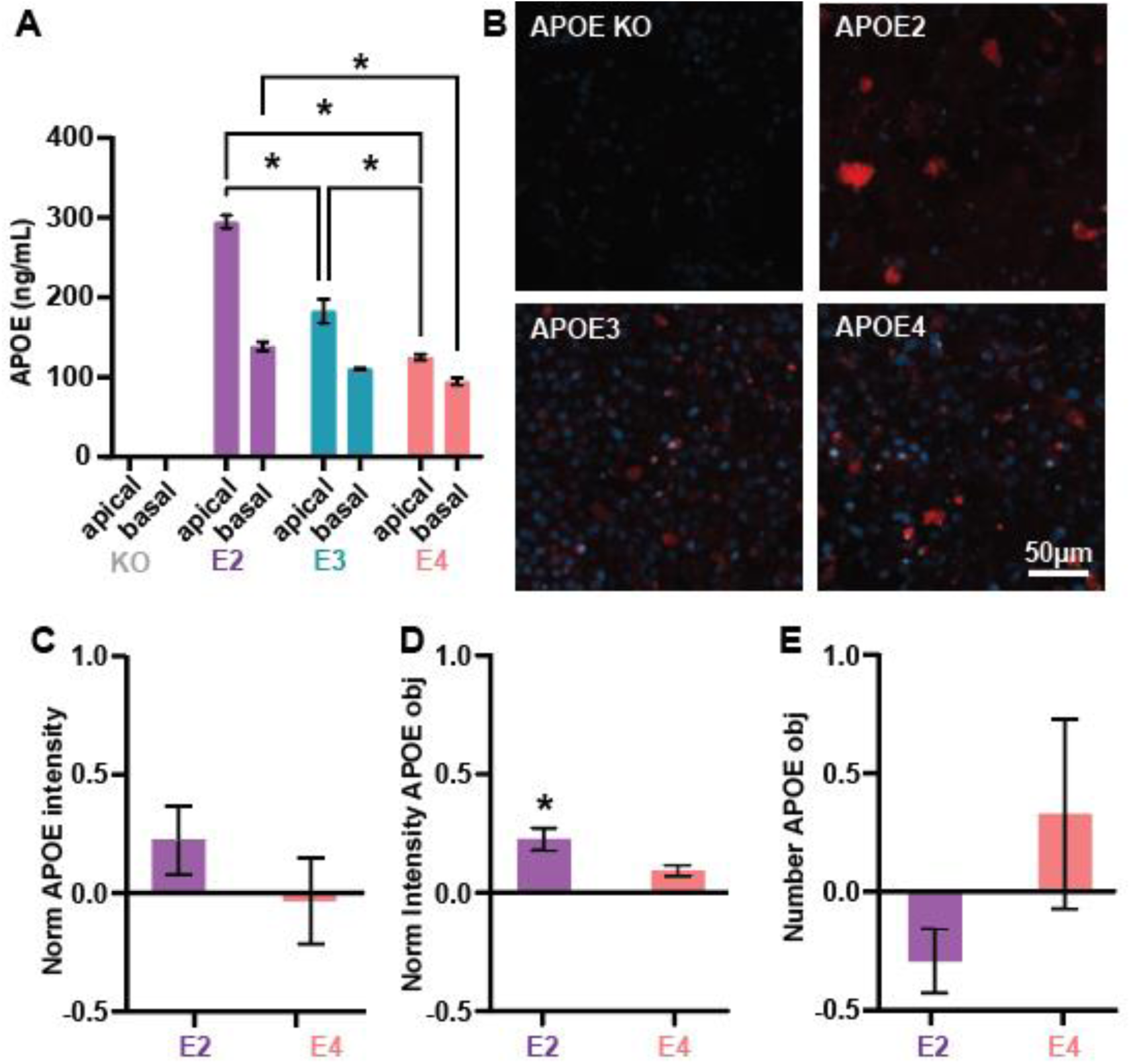
APOE expression is isoform-dependent in iPS-RPE cells. **A)** APOE concentration measured in apical and basal media supernatants of APOE iPS-RPE cells grown on transwell inserts. **B)** Representative images of APOE immunofluorescence staining in 8-week-old iPS-RPE cells. Scale bar indicates 50µm. **C)** Quantification of APOE staining intensity within APOE positive assemblies normalized to APOE3. **D)** Number of APOE positive assemblies identified in maximum intensity projections. Data normalized to APOE3. **E)** Whole-field APOE staining intensity in maximum intensity projections. Data normalized to APOE3. * used to indicate any p <0.05 for simplicity. Data are from three biological replicates.

Because some reports suggest that APOE can accumulate or aggregate within cells, we next sought to understand if intracellular APOE concentration also varied by APOE isoform. We performed immunofluorescence staining to visualize APOE in iPS-RPE cells. Staining revealed that intracellular APOE abundance is highly heterogeneous, with robust intracellular accumulation observed in a small subpopulation of cells, while APOE was low to undetectable in most iPS-RPE (**Fig 3B**). APOE staining generally appeared to be in puncta but can be observed to fill the whole cell in a small population (**Fig 3B**). While APOE IF staining was generally stronger in APOE2 samples compared to APOE3 and APOE4, data were variable and did not reach significance when comparing APOE staining intensity (**Fig 3C**). To better quantify APOE staining patterns, APOE-positive objects were segmented using Arivis Vision4D software^50^ for further analysis. Objects identified in APOE2 cultures tended to be fewer in number (E3>E4>E2, **Fig 3D**) but had significantly higher intensity staining than APOE3 or APOE4 objects (**Fig 3E**). Staining was at background levels in APOE KO iPS-RPE and no objects were segmented using our automated pipeline.

### Bulk RNAseq reveals lipid pathway dysregulation in iPS-RPE cells

Our initial observations suggested that APOE KO cells have considerable morphological deficits that could be caused by alterations to potentially many different cellular processes or pathways. Likewise, APOE2 cells show similar, but milder, morphological deficits. We took a broad profiling approach to better understand how iPS-RPE function may be altered by *APOE* loss or expression of *APOE2* and *E4* by performing bulk RNA sequencing on 4-week-old iPS-RPE cells. Using *APOE3*, the most common allele in the human population, as our reference we performed differential gene expression analysis comparing to identify genes altered in *APOE2*, *APOE4*, and *APOE* KO iPS-RPE cells. Gene expression changes are most pronounced between *APOE3* and *APOE* KO iPS-RPE cells, with over 3000 differentially expressed genes (DEGs) identified (**Fig 4A**). Changes between *APOE2*-*APOE3* and *APOE4*-*APOE3* were more subtle, with 444 and 642 DEGs identified respectively (**Fig 4A**).

**Figure 4:**
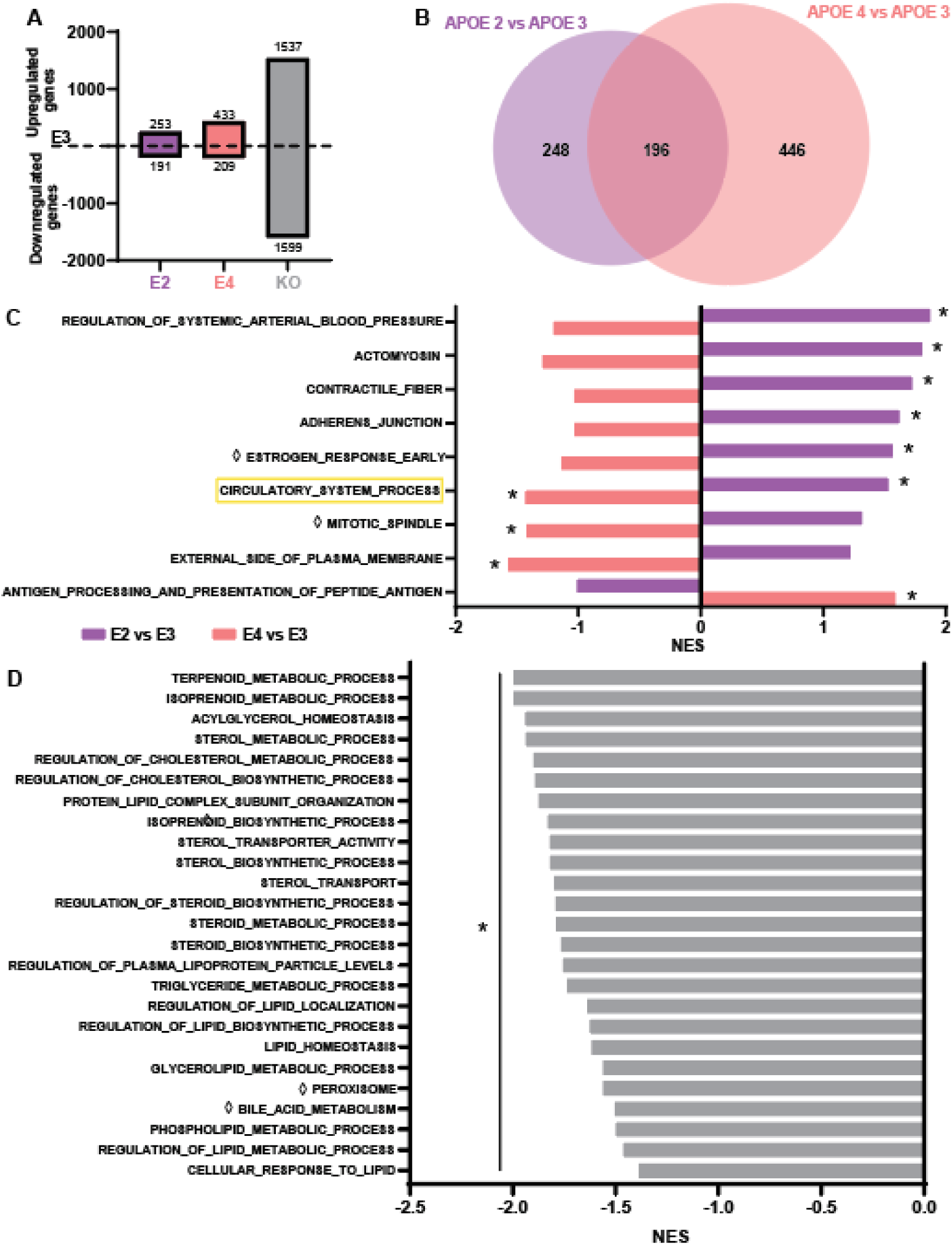
Bulk RNAseq analysis reveals lipid pathway dysregulation in APOE KO iPS-RPE. **A)** Number of up- and downregulated genes in APOE2, APOE4, and APOE KO iPS-RPE compared to APOE3. **B)** Number of DEGs (differentially expressed genes) in APOE2 and APOE4 iPS-RPE compared to APOE3. Of all DEGs, 196 were shared between APOE2 and APOE4. **C)** Oppositely regulated pathways in APOE2 and APOE4 iPS-RPE compared to APOE3. Pathways shown have NES >1 or <-1. **D)** Selected lipid-related pathways downregulated in APOE KO compared to APOE3. Pathways shown have <-1 NES and are significantly downregulated. ◊ indicates pathway is from the Hallmark Database, all others are from GO database. * used to indicate any p <0.05 for simplicity.

We reasoned that genes of high interest in AMD would be differentially regulated in both *APOE2* and *APOE4* but would likely have opposite directionality of regulation. For example, a gene of interest might be upregulated in APOE2 cells and downregulated in APOE4 cells compared to APOE3 control. Of the DEGs identified in APOE2 and APOE4 cells, only 196 showed significant changes in both APOE2 and APOE4 iPS-RPE (**Fig 4B**), representing a relatively small portion of the total genes. We next sought to identify pathways that might be impacted by expression of *APOE2* and *APOE4* in order to better understand which cellular processes could underlie AMD risk or protection in RPE cells. We identified 22 pathways that were regulated in opposite directions in APOE2 and APOE4, but only 9 had normalized enrichment scores (NES) of 1 or greater in both APOE2 and APOE4 (**Fig 4C**). Of these, only the Circulatory_System_Process gene set was significantly differentially expressed in both APOE2 and APOE4. Members of this pathway encode cholesterol transporters, lipoprotein receptors, and a diverse array of other proteins involved in vascular biology. However, of these 602 genes, only *ADAMTS16* and *NPPB* were oppositely regulated in APOE2 and APOE4 iPS-RPE cells (**Fig S2A**).

Because differences were subtle in APOE2 and APOE4, we considered the larger gene expression changes observed between APOE KO and APOE3 cells. Pathway analysis revealed 289 pathways from the GO, KEGG, and HALLMARK databases to be significantly differentially regulated in APOE KO cells compared to APOE3. While pathways spanned a diverse array of cellular processes, we observed that a considerable number of differentially regulated pathways concerned lipid metabolism and transport (**Fig 4D**). Notably, all these pathways were found to be downregulated in APOE KO cells compared to APOE3, suggesting that loss of *APOE* may have widespread impact on lipid metabolism in iPS-RPE cells.

### APOE2 impairs response to lipid challenge in iPS-RPE cells

Our findings show that APOE is important for RPE health and our bulk RNAseq results suggest that loss or dysfunction of APOE may impact multiple aspects of lipid metabolism and circulatory function. Since lipid trafficking is a key function of APOE across cell types, we chose to investigate whether APOE isoform or loss of APOE impacts lipid trafficking in iPS-RPE cells. First, we characterized whether APOE expression is responsive to a lipid challenge in iPS-RPE cells by exposing cells to high extracellular cholesterol (**Fig 5A**). Cells were treated with cholesterol-bound cyclodextrin (50µg/mL), also known as “soluble cholesterol,” for one week in typical iPS-RPE growth media, after which APOE expression was measured in media supernatants using ELISA. APOE was not detected in the APOE KO line, as expected (**Fig 5A**). We found that exposure to this lipid challenge increased APOE expression in all APOE-expressing lines (**Fig 5A**). Because APOE protein levels are higher in APOE2 media supernatants at baseline, we calculated the fold change in APOE concentration in response to cholesterol challenge. The fold increase in APOE concentration in media was 3 times greater in APOE4 cells than APOE2 cells, indicating a stronger cholesterol-mediated response in APOE4 cells (**Fig 5B**).

**Figure 5:**
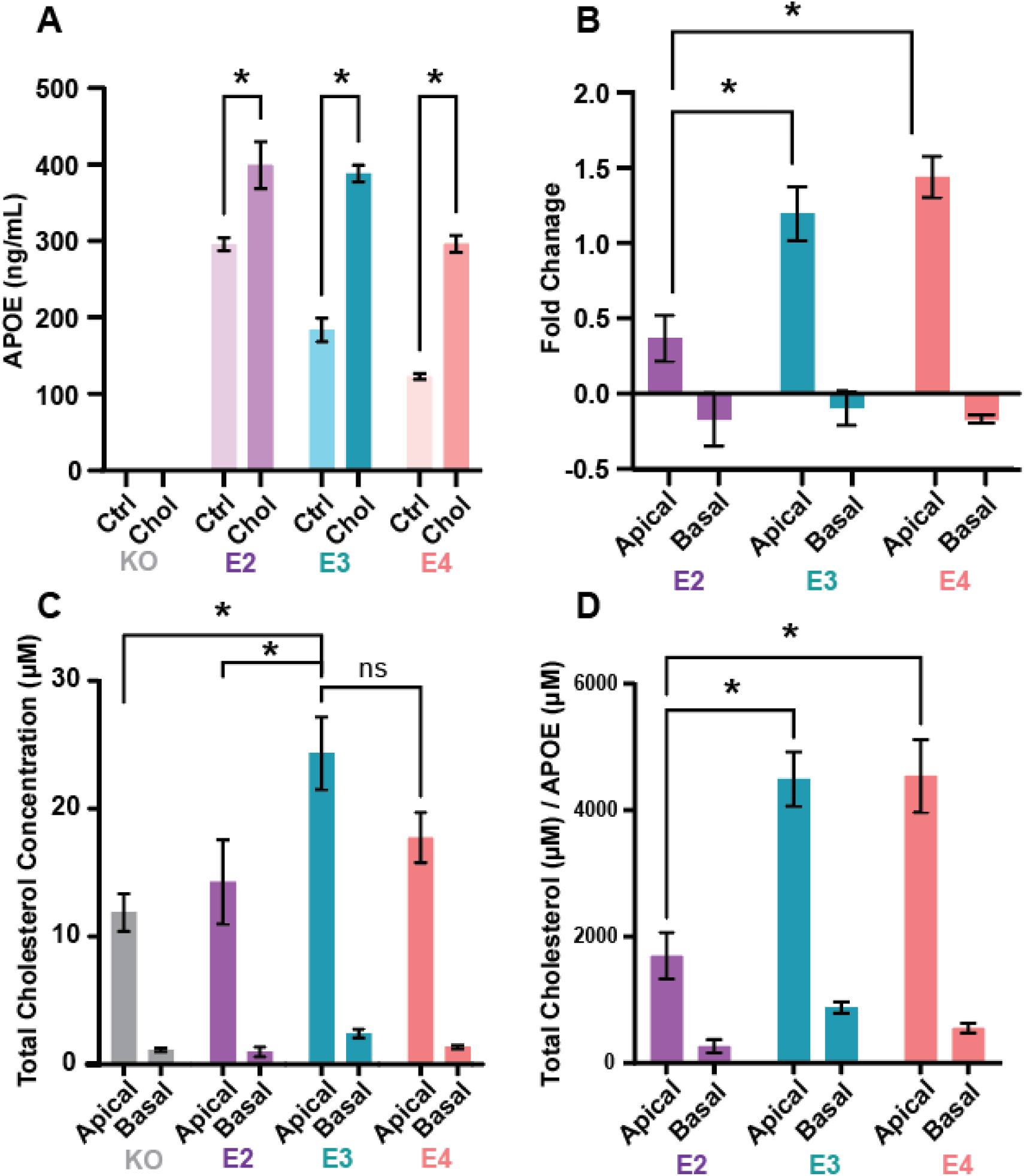
APOE isoform impacts lipid transport and response to lipid challenge. **A)** APOE expression in apical media measured by ELISA in untreated (Ctrl) and cholesterol challenged (Chol, 50µg/mL) APOE iPS-RPE. **B)** Fold Change in APOE protein expression in apical and basal media supernatants measured by ELISA in cholesterol challenged APOE iPS-RPE cells. **C)** Total cholesterol concentration in apical and basal media supernatants measured by luminescence assay in APOE iPS-RPE cells. **E)** Ratio of total cholesterol measured in media supernatants to APOE measured in the same supernatant sample. * used to indicate any p <0.05 for simplicity.

Neutral lipids are transported in the extracellular space in the core of lipoprotein particles. APOE is a prominent lipoprotein in the central nervous system and is thought to be a component of these particles in the brain and retina. However, several lipoproteins, such as APOA1 and others, may also be expressed in these tissues. To better understand whether APOE is involved in lipid trafficking in RPE cells, we measured cholesterol content in RPE-conditioned media. Cholesterol was present in RPE-conditioned media from cells of all tested genotypes (**Fig 5C**), suggesting that cholesterol efflux from RPE cells does not require APOE. Cholesterol concentration was higher in the apical compartment and lower in the basal compartment (**Fig 5C**). Interestingly, APOE KO and APOE2 cells had lower total cholesterol concentration in both apical and basal media compared to APOE3 cells (**Fig 5C**), but neither differed from APOE4. However, since APOE abundance is higher in APOE2 cells, the ratio of cholesterol to APOE is significantly decreased in APOE2 cells compared to those expressing APOE3 or APOE4 (**Fig 5D**), perhaps suggesting that the secreted APOE2 is poorly lipidated in these cells.

Intracellular lipids are frequently stored as lipid esters in lipid droplets, organelles consisting of a single phospholipid layer surrounding a core of TAGs and cholesteryl esters^51^. We hypothesized that lipid droplet abundance of morphology may be impacted by APOE isoform or loss of APOE. APOE4 expression is known to increase lipid droplet abundance compared to APOE3 in CNS cell types, such as astrocytes^52–55^. Lipid droplets were visualized using a lipophilic dye (BODIPY 493-503) in APOE iPS-RPE cells in control media or media supplemented with 50ug/mL soluble cholesterol for seven days. Due to differences in staining intensity in different experiments, results are presented normalized to APOE KO. BODIPY 493/503 staining intensity was lower in APOE2 and APOE4 cells compared to APOE KO cells, but slightly higher in APOE3 cells (**Fig 6A-B**). Individual lipid droplets were segmented using Arivis Vision4D software, revealing a similar pattern of results (data not shown). Treatment with cholesterol-enriched media did not change BODIPY staining intensity in APOE2, APOE3, and APOE KO cells, but greatly increased lipid droplet accumulation in APOE4 cells (**Fig 6A, 6C**). Greater intake and storage of neutral lipids in APOE4 iPS-RPE could facilitate metabolism of excess lipids for energy or degradation of excess lipids, such as by lipophagy. Taken together, APOE4 cells show a greater response magnitude to a lipid challenge, both in APOE protein expression and lipid storage.

**Figure 6:**
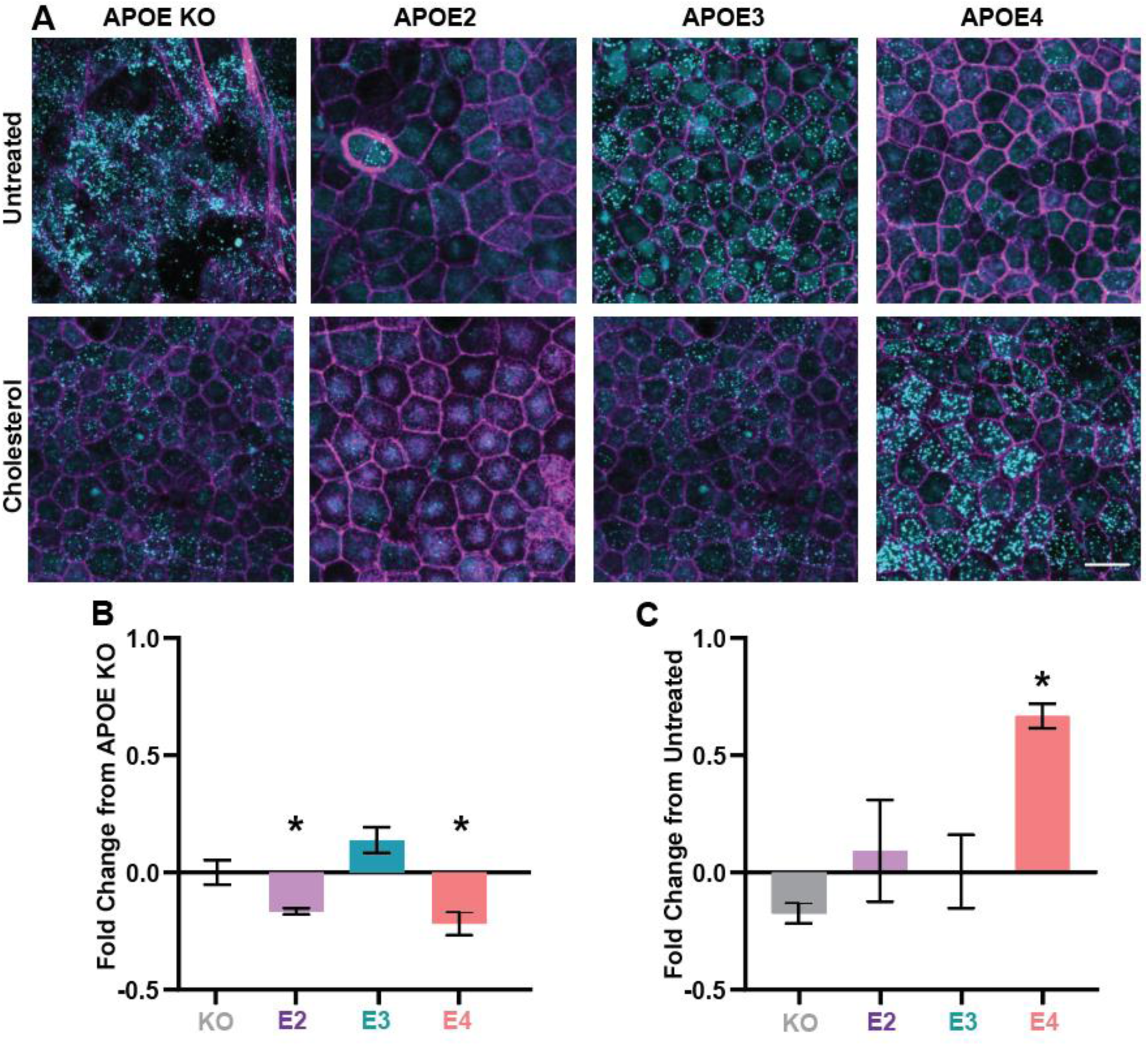
APOE4 increases neutral lipid storage in response to cholesterol challenge in iPS-RPE. **A)** Representative images of APOE iPS-RPE cells grown in typical growth media (Control) or subjected to a cholesterol challenge (Cholesterol) for 7 days. Blue = DAPI (nuclei), Red = Phalloidin (β-Actin), Green = BODIPY (neutral lipid). Scale bar indicates 20µm. **B)** Change in total BODIPY fluorescence normalized to APOE KO in untreated iPS-RPE cells. **C)** Change in BOIDPY signal in cholesterol challenged iPS-RPE compared to untreated iPS-RPE cells. * used to indicate any p <0.05 for simplicity.

## Discussion

In the current work, we have expanded the knowledge of how APOE functions in iPS-RPE cells. Loss of APOE results in abnormal RPE morphology, including large cell size and aberrant pigmentation. APOE KO iPS-RPE cells also show poor barrier function that deteriorates over time as cells are lost. Furthermore, loss of APOE alters gene expression in iPS-RPE cells, with especially strong impact on a number of lipid synthesis, metabolism, and transport pathways. We observed more subtle differences between APOE2 and APOE4 iPS-RPE, as might be expected from variants that typically only alter risk for disease or disease progression at advanced patient ages. For example, changes in gene expression between the two variants were relatively limited, though several differentially regulated genes and pathways have been identified here to be explored in future studies. Across many *in vitro* and *in vivo* systems, an increased abundance of APOE2 relative to APOE3 and APOE4 has been observed. We find that this is also true in iPS-RPE cells, with secreted APOE2 being almost twice as abundant as APOE4 in our culture system. Perhaps the most striking difference we observe is the strong response of APOE4 iPS-RPE cells to a lipid challenge. APOE4 iPS-RPE strongly upregulate APOE in response to high extracellular cholesterol and may be able to transport lipids more efficiently by forming particles with a higher APOE:Lipid ratio. Lastly, APOE4 iPS-RPE show a unique response to lipid challenge by storing excess lipids in lipid droplets, which could possibly protect these excess lipids oxidative damage.

The ultimate goal of our work presented here is to better understand how APOE might contribute to the development of AMD. While much work remains to be done to fully understand why APOE2 increases risk for AMD and APOE4 is protective, our studies have revealed several interesting leads to be followed up in future work. One significant observation is that APOE2 iPS-RPE show some shared characteristics with APOE KO iPS-RPE, possibly suggesting that APOE2 has reduced functionality in RPE cells, despite having high levels of expression. Like APOE KO cells, APOE2 cells show increased size relative to APOE3 and APOE4 cells. Furthermore, APOE2 cells showed equivalent cholesterol transport capacity as APOE KO cells, further suggesting that APOE2 may not function adequately in lipid transport in RPE cells. In contrast, APOE2 cells had normal barrier function, suggesting that APOE2 does not fully phenocopy loss of APOE and retains some functionality. This is consistent with observations that humans carrying the APOE2 variant do not experience overt early degeneration or RPE dysfunction and only show symptoms later in life.

In contrast, previous reports suggest that the APOE4 variant offers some protection against AMD in the human population ^6,10^. Our work suggests that iPS-RPE cells expressing APOE4 have three key differences from RPE cells expressing other APOE isoforms. Secreted APOE expression is lowest in APOE4 iPS-RPE cells, consistent with findings in the CNS and systemic circulation^43,44^. While lower APOE concentrations could predispose patients to diseases in these systems, we know that APOE is an abundant protein component of drusen in AMD and has been known to enhance the stability of protein aggregates in other systems^30,33,56–60^. Less secreted APOE may be an advantage in the eye, since fewer unlipidated APOE or APOE-bound lipoproteins would be available to contribute to drusen formation and resulting inflammatory response. We also find that APOE4 iPS-RPE increase APOE expression more strongly than cells carrying the other isoforms in response to a lipid challenge. Taken together, APOE4 cells have low APOE expression at baseline, but are able to upregulate APOE secretion to mobilize excess lipids efficiently when extracellular lipid concentrations are high, such as in AMD. Lastly, we find that cells expressing APOE4, unlike any of the other tested variants, increase the amount of neutral lipids stored in lipid droplets when faced with a lipid challenge. While further work will be needed to fully understand the adaptive advantage of this proliferation of lipid droplets, it is known that lipid droplets can be a privileged compartment to protect lipids from oxidation and provide an energy store for cells^51^. Previous work has shown that RPE cells are sensitive to oxidative damage^61,62^ and lipid-oxidation-dependent forms of cell death, such as ferroptosis^63–68^. Sequestering vulnerable lipids in droplets could help to mitigate oxidative lipid damage in RPE cells.

Our study expands the current understanding of how RPE-derived APOE may impact RPE health and function. However, more work is required to understand how APOE functions in AMD and leverage this knowledge to meet patient needs. One major outstanding question raised by the current work is how APOE isoforms impact gene expression in RPE cells. We found that iPS-RPE expressing either the protective *APOE4* or risk *APOE2* allele showed relatively subtle changes in gene expression compared to iPS-RPE expressing *APOE3*. This is not surprising, since we would expect that changes in cellular function must be relatively subtle if carriers of these variants have normal function for decades of life before being at risk to develop age-related diseases such as AMD or Alzheimer’s disease. However, our initial assumption was that we would be able to identify key pathways and genes that are significantly regulated in opposite directions in APOE2 an APOE4 cells; for example, “pathway X” might be enriched in APOE2 cells and de-enriched in APOE4 cells. We were only able to identify one pathway meeting this description, which may prove to be of high relevance since it pertains to circulatory system function, a known vulnerability in AMD. In contrast, we identified many pathways that were regulated in the same direction in APOE2 and APOE4 cells, but with vastly different magnitudes of effect. Interestingly, APOE4 tended to have the larger effect size, with APOE2 having less impact. This has led us to hypothesize that APOE2 and APOE4 may not confer risk and protection respectively by imposing opposite influences on cellular function. Instead, we can conceptualize a system in which APOE2 may alter pathway function slightly, leading to mild cellular dysfunction, but APOE4 greatly alters functioning in the same pathway, triggering compensatory changes in other cellular functions, ultimately leading to protection.

Our work also does not consider the role of APOE and APOE lipoprotein particles derived from other cell types in the eye. Indeed, RPE cells highly express APOE receptors, suggesting that the RPE is a key site of action for APOE function in the eye. However, we also know from previously published single nucleus RNAseq data^69,70^ that RPE cells express less *APOE* compared to cells in the neural retina, such as Müller glia, astrocytes, and immune cells (**Fig S2C-D**). It remains unexplored whether APOE lipoprotein particles sourced from cells in the neural retina and the RPE/choroid have different physical characteristics or lipid compositions that could impact their function in lipid transport or cellular signaling. In fact, the majority of APOE is secreted apically in our *in vitro* system, yet drusen are located on the basal side of the RPE, suggesting that APOE and APOE-containing lipoproteins may be sourced from systemic circulation as well as the retina *in vivo*. It is also unknown whether other cell types show the dynamic regulation of APOE expression in response to lipid stressors that we observed in RPE cells and how this could be impacted by the disease state, especially if they are physically removed from degenerating RPE in AMD. Future work to characterize APOE lipoprotein particles derived from different cell types in the neural retinal and RPE/choroid could clarify APOE function in the eye. More complex culture systems, such as co-culture of APOE-secreting Müller glia with RPE cells, or treating RPE cells with Müller glia-derived APOE lipoprotein particles^71^, could illuminate cell-cell interactions in normal function and disease.

The work presented here establishes new findings that relate APOE function in RPE cells to possible AMD-relevant dysfunction. We have shown that APOE is required for continued RPE health in *in vitro* systems, and that the AMD-risk APOE2 allele may lead to delayed or altered RPE maturation. Furthermore, we have shown that APOE4 iPS-RPE cells are able to dynamically regulate APOE expression and lipid storage in response to lipid challenge, perhaps suggesting an advantage in AMD, which is characterized by large lipid deposits in the form of drusen. At the same time, the current work opens many new lines of investigation, such as how multiple APOE-secreting cell types may work together in the multicellular *in vivo* context and how changes to lipid storage may confer protection from AMD. Future work addressing these outstanding questions will better clarify how APOE variants confer risk and protection in AMD and could potentially aid the development of effective treatments for AMD patients.

## Methods

### Human Tissue Use

The Lions Eye Institute for Transplant & Research (LEITR; Tampa FL) provided postmortem human eye tissue with consent of donors or donors’ next of kin. Donations were collected in accordance with the Eye Bank Association of America (EBAA) medical standards, country (United States) and state (Florida) law for human tissue donation, the Declaration of Helsinki and FDA regulations, and Novartis guidelines regarding research using human tissues. All experimental procedures were performed in accordance with Novartis policies for research using human tissues.

### Post-Mortem Human Donor Eye Collection and Histology

All study tissue samples were preserved within 6 hours post-mortem or less. Donor eyes for histology were injected in the vitreous with 100 µl of Modified Davidson’s Fixative (MDF, H0290-500ML) and preserved as whole globes in that same fixative. Eyes were fixed for 48 hours and transferred to 70% alcohol for an additional 48 hours, all at room temperature. After fixation, globes were sectioned horizontally so that the sections included the optic nerve head and the macula. Five-micron sections were dried at 60°C for 24 hours prior to immunohistochemistry (IHC), and hematoxylin and eosin staining (H&E). Procedures conducted as previously described ^45^.

### Development of APOE Cells

Human iPSCs expressing APOE2, APOE3, and APOE4 or APOE KO were obtained from Alstem. Briefly, human iPSCs homozygous for APOE4 (APOE4/APOE, Alstem IPS16) were genetically modified using CRISPR to generate lines homozygous for APOE2 and APOE3 (Alstem lines IPS46 and IPS26). APOE KO cells were generated by inducing a stable homozygous 100 bp deletion in the exon 2 of APOE gene. All edits were verified using Sanger sequencing. Stem cell differentiation protocol can be found on the Alstem website. Stem cells were differentiated to Retinal Pigment Epithelium cells (iPS-RPE) by Alstem. Briefly, iPSCs were grown on Matrigel coated dishes until they reached 80% confluence. Culture media was switched DMEM high glucose with 1x Nonessential Amino Acids and 20% knockout serum replacement (KSR). To this media, 10 nM Nicotinamide was added during Day (D)0-D7, 100 ng/mL Activin A was added D7-D14, and CHIR99021 was added D14-D21. Following D21, cells were moved to 4% KSR differentiation media for expansion. Following expansion, cells were cryopreserved and shipped to our site for use. Alstem does not provide item numbers for the exact differentiation reagents in their materials.

### Cell Culture

Commercially available iCell iPS-RPE (R1102) cells were sourced from FujiFilm/Cellular Dynamics (Donor #01279). Our other control cell line, iPS-RPE 01434 was a custom line developed by FujiFilm/Cellular dynamics for our group. Donor information can be found on the FujiFilm/Cellular Dynamics website.

iPS-RPE cells were thawed into Lonza RPE media (Lonza 00195406) +2% Fetal Bovine Serum (Gibco 10082147) and anti-mycotic/anti-microbial (Corning 30-004-CI) and seeded directly into the culture vessel required for each experiment (e.g. 24 well transwell plate, 96 well plate, etc.). Cells were maintained in this medium for two weeks with complete media changes every 2-3 days. After two weeks, cells are maintained in Lonza X-VIVO 10 (Lonza, BP04-743Q) with anti-mycotic/anti-microbial (Corning 30-004-CI) with media changes every 2-3 days. Cells were matured for 4-8 weeks before use in experiments as specified in the manuscript.

For lipid challenge experiments, growth media was supplemented with soluble cholesterol (50 µg/mL, C4951) for 72 hours or 1 week. Media changes were completed every 2-3 days.

Transepithelial electrical resistance (TEER) was measured using a Millicell ERS-2 (Millipore) manual epithelial volt-ohm meter. Before use, the probe was sanitized in 70% ethanol for 15 minutes, then washed twice in sterile DPBS. Measurements were recorded manually and normalized to the total surface area of the culture surface, resulting in measurements expressed as Ω/cm^2^.

### Cholesterol Assay

Mature iPS-RPE cells were grown as described above in 24W plates with transwell inserts (Falcon 353095). Apical and basal media was collected from cells growth in maintenance media or media supplemented with cholesterol (50 µg/mL; Sigma-Aldrich C4951). Media samples were spun for 3 minutes at 1000 g in a temperature-controlled centrifuge (4 °C) to pellet any floating cells or cellular debris that might lead to erroneous signal in the assay. Media samples were used immediately in the Cholesterol/Cholesterol Ester-Glo (Promega, J3191). Assays were performed according to manufacturer instructions. For the cholesterol assay, samples were read with and without addition of Cholesteryl Esterase to enable measurement of free, esterified, and total cholesterol. Luminescence was read on a Tecan Spectrometer. Data analysis was performed in GraphPad Prism.

### Immunofluorescence and Staining

Cells were fixed in 4% paraformaldehyde (Thermo, 043368.9M) in the apical and basal chamber for 15min at RT followed by 3 washes with PBS. Plates were sealed with parafilm to prevent evaporation and stored at 4 °C until use.

For immunofluorescence, transwell membranes were permeabilized with 0.02% triton X-100 in PBS for 10 minutes at RT with gentle shaking, followed by 3 washes with PBS. Membranes were blocked with 1% BSA in PBS (VWR, 76286-966) in both apical and basal chambers for 1 hour at RT with gentle shaking. Primary antibodies for APOE (1:500; R&D Systems, MAB41441), ZO-1 (1:200; Invitrogen 33-9106) and CRALBP (1:500, Abcam, ab243664) were added overnight in blocking buffer at 4 °C with gentle shaking. Plates were washed 3 times with PBS and secondary antibodies were applied in blocking buffer for 1 hour at RT with gentle shaking: anti-rat 647 (1:500; Jackson Immuno, 102647-386), anti-rabbit 594 (1:500; Jackson Immuno, 102645-414), anti-mouse 488 (1:500; Jackson Immuno 102650-156). Hoechst nuclear stain (1:1000, Thermo 62249) was added with secondary antibodies. Membranes were washed three times with PBS. Membranes were removed from the transwell scaffold and mounted on slides with SlowFade mounting media (Invitrogen, S36972) and covered with a cover slip. Slides dried overnight and edges were sealed with clear paint.

For staining, dyes were applied to transwells in the apical and basal chambers in PBS and incubated for 1 hour at RT with gentle shaking using the following dilutions: BODIPY 493/503 (1:500, item), Phalloidin 488 or 647 (0.15µM, Invitrogen 12319 or A30107), filipin (500µg/mL, SAE0087). Dyes were co-applied with a nuclear stain, either Hoechst (1:1000, Thermo 62249), Sytox 488 (1:5000; Invitrogen, S7020), or Sytox Deep Red (1:5000; Invitrogen, S11381). Dye incubation was followed by three washes in PBS with gentle shaking. Membranes were removed from the transwell scaffold and mounted on slides with SlowFade mounting media (Invitrogen, S36972) and covered with a cover slip. Slides dried overnight and edges were sealed with clear paint.

All membranes were imaged using a Zeiss 990 AiryScan confocal microscope at 20X. Four sites were imaged on each membrane and these four observations were averaged to generate one experimental observation. Three or more independent membranes were images for each condition. Data analysis was performed using custom automated image analysis pipelines developed in Arivis (Zeiss) as well as ImageJ/Fiji (NIH) and Zen Blue (Zeiss). Further calculations and data analysis were performed in GraphPad Prism 9.

### ELISA

Secreted APOE was measured using an in-house developed ELISA. Nunc Maxisorp 96W plates (Thermo, 442404) were coated with rat anti-APOE capture antibody (Invitrogen, 701241) diluted 1:1000 in PPS overnight with gentle shaking. Plates were washed 4 times with 300 uL of TBST wash buffer. Media supernatant samples were added to each well (100 uL) along with APOE standard (Peprotech, 350-02) in concentrations ranging from 5 µg/mL to 1.2 ng/mL in 1:4 dilution steps in identical media to the samples, along with a blank media sample. Samples were incubated for 1 hour at RT with gentle shaking. Plates were washed 4 times with 300 uL of TBST wash buffer. Detection antibody (R&D Systems, MAB41441) diluted 1:1000 in blocking buffer (VWR, 76286-966) was added for 1 hour at RT. Plates were washed 4 items with 300 uL of TBST wash buffer. HRP-conjugated secondary antibody (Invitrogen, A18745) was added 1:2000 in blocking buffer for 1 hour at RT with gentle shaking. Plates were washed 4 times with 300 uL of TBST buffer. TMB substrate (Thermo 34028) was added for approximately 5 minutes and reaction was stopped with sulfuric acid Stop solution (Invitrogen, SS04). Absorbance was immediately read at 450 nm. Total protein concentration was measured (BioRad, 220115-411) and used for normalization.

### RNA Isolation

Total RNA was isolated from 4-week-old iPS RPE cells expressing APOE2, APOE3, APOE4 or APOE KO. Cultures were maintained using standard culture procedures outlined above. RNA isolation was performed using the Zymo Tri-reagent kit (R2050) as per manufacturer instructions with some minor alterations. Following lysis with 300 uL of Tri-reagent and addition of 300 uL of ethanol, samples were spun at 1000 g for 1 minute to pellet unlysed cellular debris common in RPE cell samples. The supernatant was removed from the pelleted debris and used for the remainder of the protocol. The optional DNase step was included. RNA concentration was measured using a NanoDrop instrument and concentration and RIN value was measured using an Agilent 2100 Bioanalyzer. Samples stored frozen at −80 °C until shipment to Genesupport (Plan-Les-Ouates, Switzerland) for sequencing.

### RNA Sequencing

Two hundred fifty nanograms of total RNA per sample were used as input to the Illumina TruSeq Stranded mRNA Library Prep Kit (reference 20020595). The libraries were generated per manufacturer’s specifications, except Illumina adapters provided with the kit were replaced with NEXTFLEX® Unique Dual Index Barcodes (Perkin Elmer, references NOVA-514150, NOVA-514151, NOVA-514152, NOVA-514152). RNA-Seq libraries were amplified using the Illumina Truseq primer mix (14 cycles). The PCR amplified RNA-Seq library products were then quantified using the Quant-iT™ dsDNA Assay Kit, High Sensitivity (In Vitrogen, reference Q33120) on a Gemini XPS (Molecular Devices, reference XPS05542). The library profiles were checked using High Sensitivity NGS Fragment Analysis kit (1-6000bp) (Agilent Technologies, reference DNF-474-1000) on the 5300 Fragment Analyzer System. The libraries were diluted to 5 nanomolar in Elution Buffer, pooled, denatured, and loaded at 300 picomolar on Illumina NovaSeq 6000 flowcells S1 and SP NovaSeq 6000 Reagent Kit v1.5. The RNA-Seq libraries were sequenced 50 base pair paired-end with 8 base dual indexes. The sequence intensity files were generated on-instrument using the Illumina Real Time Analysis software. Paired-end 50 bp reads were generated for an average of 34 million read pairs per sample. FASTQ files were extracted using Illumina’s bcl2fastq software (v2.17.1.14).

### Data Analysis

For all common laboratory experiments, data were analyzed using Microsoft Excel and GraphPad Prism 9. Data were organized and, in some cases, normalized using Excel. All statistical analysis and curve fitting was performed in Prism.

For bulk RNA sequencing, we used the Exon Quantification Pipeline [1] (version 2.5) with STAR ^72^ (version 2.7.3a) as the alignment tool to align the reads against the human genome reference files from Ensembl version 98 ^73^ and quantify gene expression. The normalization and differential expression (DE) analysis were performed using edgeR and limma-voom R packages to identify the DE genes (absolute log 2 fold change ≥1 and adjusted p value <0.05). The enriched pathways were obtained using GSEA.

## Author Contributions

Sarah E.V. Richards – conception and design of work; acquisition, analysis, and interpretation of data; drafted the work

John Demirs – acquisition of data; drafted section of the work

Sandra Jose – analysis of data; drafted section of the work

Lin Fan – acquisition of data; design of work

YongYao Xu – acquisition of resources (cell lines)

Robert Esterberg – acquisition of resources (cell lines)

Chia-Ling Huang – analysis of data

Christopher W. Wilson – design of the work; approval of the work

Magali Saint-Geniez – conception of the work; interpretation of data; approval of the work

Sha-Mei Liao – conceptualization and design of the work; acquisition, analysis, and interpretation of data

---

**Figure S1:**
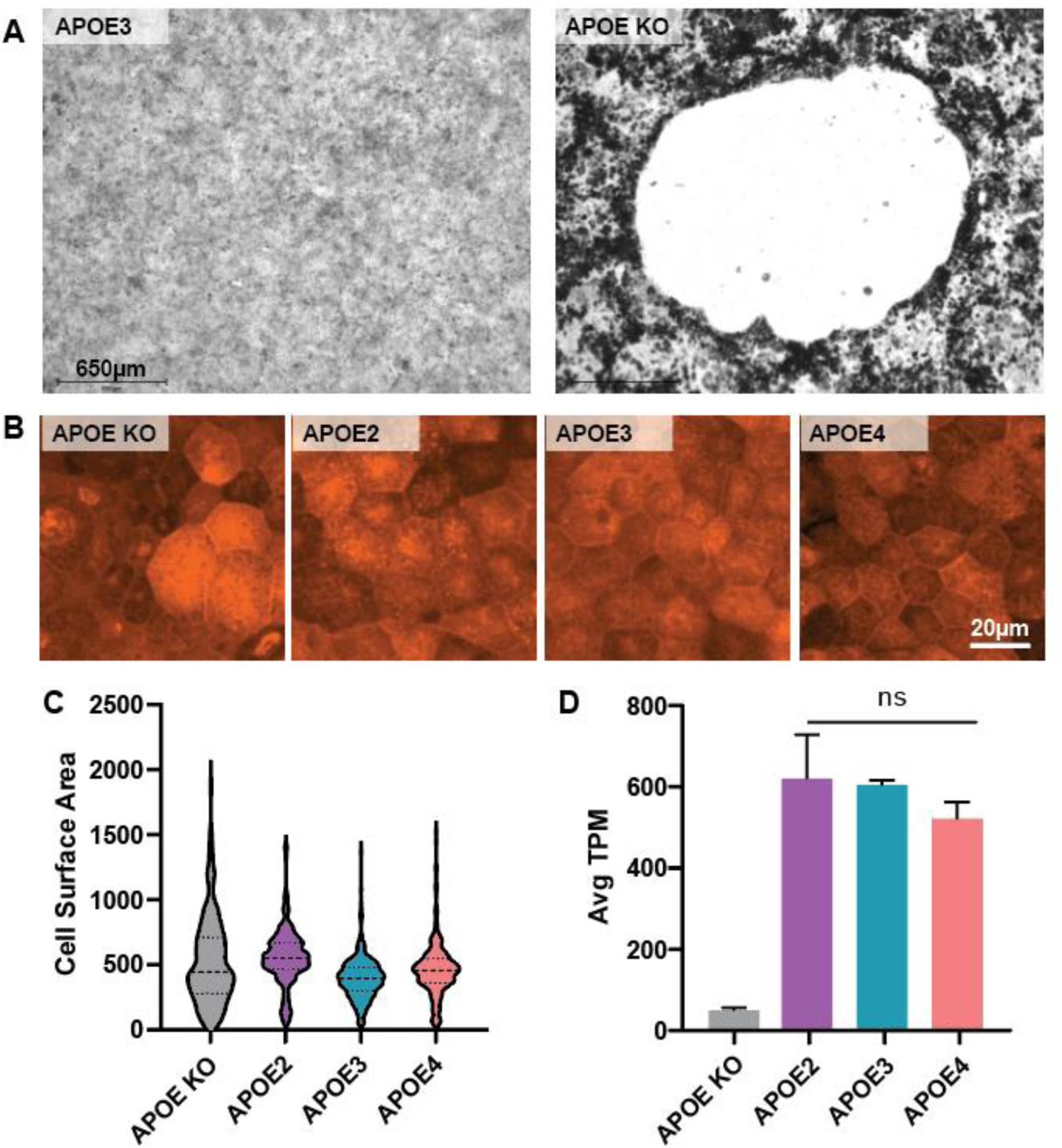
Additional characterization of *APOE* iPS-RPE cells. **A)** Example images of APOE3 (left) and APOE KO (right) iPS-RPE cells. Scale bar indicates 650µm. **B)** Example images of CRALBP staining in 8wk old APOE KO, APOE2, APOE3, and APOE4 iPS-RPE cells. Scale bar indicates 20µm. **C)**Violin plots of cell surface area (µm^2^). Heavy dashed line indicates median. Light dashed lines indicate upper and lower quartiles. **D)** Average APOE transcripts per million (TPM) detected in 4wk old APOE iPS-RPE. Data selected from bulk RNA sequencing described in Figure 4.

**Figure S2:**
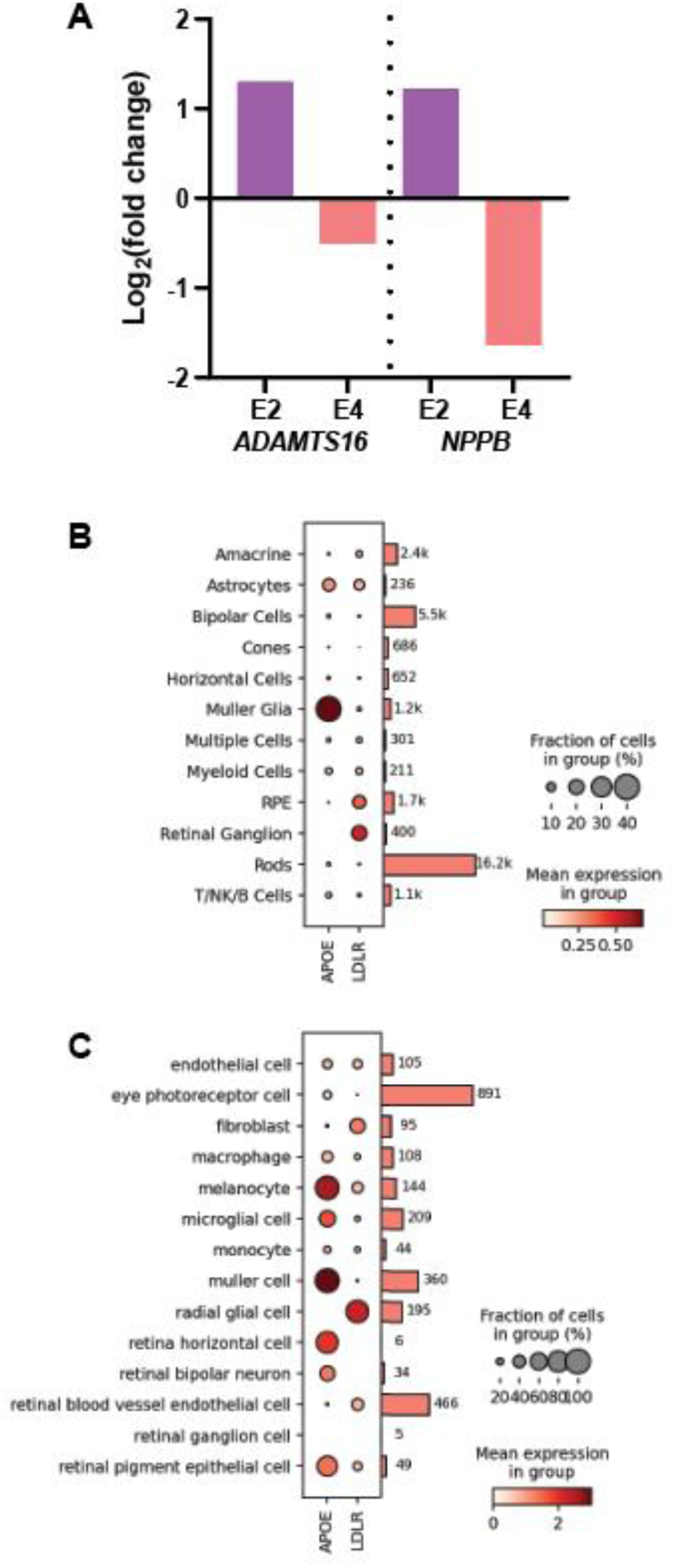
Oppositely regulated genes in APOE2 and APOE4 iPS-RPE. **A)** Log_2_ (fold change) of ADAMTS16 and NPPB in 4wk old iPS-RPE cells expressing APOE2 (violet) and APOE4 (pink). **B)** Expression of APOE and LDLR, a prominent APOE receptor in healthy aged human ocular tissue. Data replotted from Orozco et al. 2023 (https://doi.org/10.1016/j.xgen.2023.100302). **C)** Expression of APOE and LDLR, a prominent APOE receptor in healthy human ocular tissue. Data replotted from Tabula Sapiens Consortium (Chan Zuckerberg Biohub San Francisco, czbiohub.org).

## Notes

https://czbiohub.org

https://doi.org/10.1016/j.xgen.2023.100302

### Competing Interest Statement

All authors are current or former employees of Novartis Biomedical Research

